# *In vivo* structure of the *Legionella* type II secretion system by electron cryotomography

**DOI:** 10.1101/525063

**Authors:** Debnath Ghosal, Ki Woo Kim, Huaixin Zheng, Mohammed Kaplan, Joseph P. Vogel, Nicholas P. Cianciotto, Grant J. Jensen

## Abstract

The type II secretion system (T2SS) is a multi-protein envelope-spanning assembly that translocates a wide range of virulence factors, enzymes and effectors through the outer membrane (OM) of many Gram-negative bacteria. Here, using electron cryotomography and subtomogram averaging methods, we present the first *in situ* structure of an intact T2SS, imaged within the human pathogen *Legionella pneumophila.* Although the T2SS has only limited sequence and component homology with the evolutionarily-related Type IV pilus (T4P) system, we show that their overall architectures are remarkably similar. Despite similarities, there are also differences, including for instance that the T2SS-ATPase complex is usually present but disengaged from the inner membrane, the T2SS has a much longer periplasmic vestibule, and it has a short-lived flexible pseudopilus. Placing atomic models of the components into our ECT map produced a complete architectural model of the intact T2SS that provides new insights into the structure and function of its components, its position within the cell envelope, and the interactions between its different subcomplexes. Overall, these structural results strongly support the piston model for substrate extrusion.

## Introduction

The T2SS is one of seven known protein secretion systems that are used by Gram-negative bacteria to deliver proteins (effectors) across their double-membraned envelope and into the extracellular environment or into host cells^1,2^. Evolutionarily-related to bacterial type IV pili^3,4^, canonical T2SS are mainly distributed among genera of *Proteobacteria*, including representatives within α-, β-, γ-, and δ-proteobacteria^2,5,6^. However, related systems may also exist within *Spirochaetes* and *Chlamydiae*, and several other phyla^6^. The T2SS is functional in many species that are pathogenic for humans, animals, and/or plants, secreting a broad range of toxins, degradative enzymes, and other effectors^6–10^. Additionally, the T2SS is operative in a number of nonpathogenic, environmental bacteria and in some instances helps mediate symbiotic relationships^6,11^. Thus, understanding the structure and function of the T2SS is crucial, potentially assisting in the development of new strategies for battling bacterial infection^12^.

The T2SS apparatus generally employs 12 “core” components, which, for simplicity, will be referred to here as T2S C, D, E, F, G, H, I, J, K, L, M, and O ^6,13^. These components have been described as being part of four subcomplexes^6,13^. The first subcomplex is a multimer of the T2S D protein that provides the ultimate portal or gate for substrate transit across the outer membrane (OM) and out of the cell; i.e., the so-called secretin^14^. Interacting with the secretin to create a periplasm-spanning channel is an inner membrane (IM)-anchored subcomplex or platform that is comprised of T2S F, L, and M^15^. The OM-associated subcomplex and the IM-associated subcomplexes are coupled by the IM-associated “clamp protein” T2S C^15^. The third subassemblage is a pseudopilus that consists of the major pseudopilin T2S G and minor pseudopilins T2S H, I, J, and K^15^. The pseudopilus is thought to span the periplasm within the channel created by the interaction of the IM platform with the secretin and is believed to act in a “piston-” or “screw-like” fashion to drive substrate through the OM portal^6,16,17^. The fourth subcomplex is a hexamer of T2S E, a cytoplasmic ATPase that is recruited to the IM in order to “power” the secretion process^15^. The T2S O protein is an IM peptidase that processes the pseudopilins prior to their assembly into the pseudopilus^6,13^.

Structural studies of purified components and subcomplexes using crystallography, electron microscopy (EM) and nuclear magnetic resonance spectroscopy (NMR) have significantly improved our understanding of the T2SS. Some of the notable contributions include high-resolution structures of several secretins (T2S D) revealing that secretin exists as a pentadecamer^18–22^, crystal structures of different conformations of the T2SS ATPase T2S E^23,24^, a co-crystal structure of the periplasmic domains of T2S C and D^25^, a co-crystal structure of the T2S I, J, K minor pseudopilin complex^26^, a cryoEM structure of the major pseudopilin polymer^27^ and crystal structures of the soluble domains of T2S L and M^28,29^. Despite these advances, it is still not clear how the different components of the T2SS are positioned and interact with each other within an intact cell envelope *(in situ).* In recent years, electron cryotomography (ECT) has emerged as a powerful tool for studying dynamic multi-protein molecular machines *in situ* in their near native state^30^. Here, we used ECT and subtomogram averaging to reveal the first *in situ* structure of a complete T2SS. We imaged the T2SS in *Legionella pneumophila*, a pathogenic bacterium for which the biological significance of T2SS and its substrates has been extensively documented, including roles in sliding motility, lung disease, intracellular infection of macrophages and amoebae, and suppression of innate immunity ^31–37^.

## Results

### Structural details of the *L. pneumophila* T2SS *in situ*

To reveal the intact structure of the bacterial T2SS *in situ*, we used ECT to image nearly 2000 frozen-hydrated *L. pneumophila* cells (Supplementary Table 1). In our tomograms, we observed multiple electron dense “hour-glass” shaped particles (Fig. 1A-D), reminiscent of the secretin structure^14^, in the periplasm. These structures were primarily localized in the vicinity of cell poles and were not associated with any exocellular filaments (Fig. 1A, B). We also observed several top views of these particles with a diameter ~10 nm (Fig. 1E, F). While *L. pneumophila* cells encode for a Type IVa pilus (T4aP) system, which also has a secretin^38^, several lines of evidence suggest that all the structures we saw were T2SSs. First, in our ~2000 tomograms, we did not see any T4aP filaments coming out of the *L. pneumophila* cells, suggesting that under our growth conditions, T4aP is not expressed. This is consistent with earlier observation in which 30 °C was found to be better than 37 °C for the assembly of the T4aP systems^38^ (our cells were grown at 37 °C). Second, no particles were visible in 38 tomograms of a strain lacking *L. pneumophila* T2SS components T2S DE (Fig. 1G), but they were still present in a strain lacking the T4aP secretin (ΔpilQ) (Fig. 1H). In a double deletion strain lacking the T2SS secretin as well as the T4aP secretin *(ΔlspDE, ΔpilQ)*, again no particles were visible (Fig. 1I, Supplementary Table 1). Finally, our analysis of the T2SS secretin revealed distinct features as compared to the T4aP secretin (see below), confirming that the “hour-glass” shaped particles are *L. pneumophila* T2SSs.

**Fig. 1.**
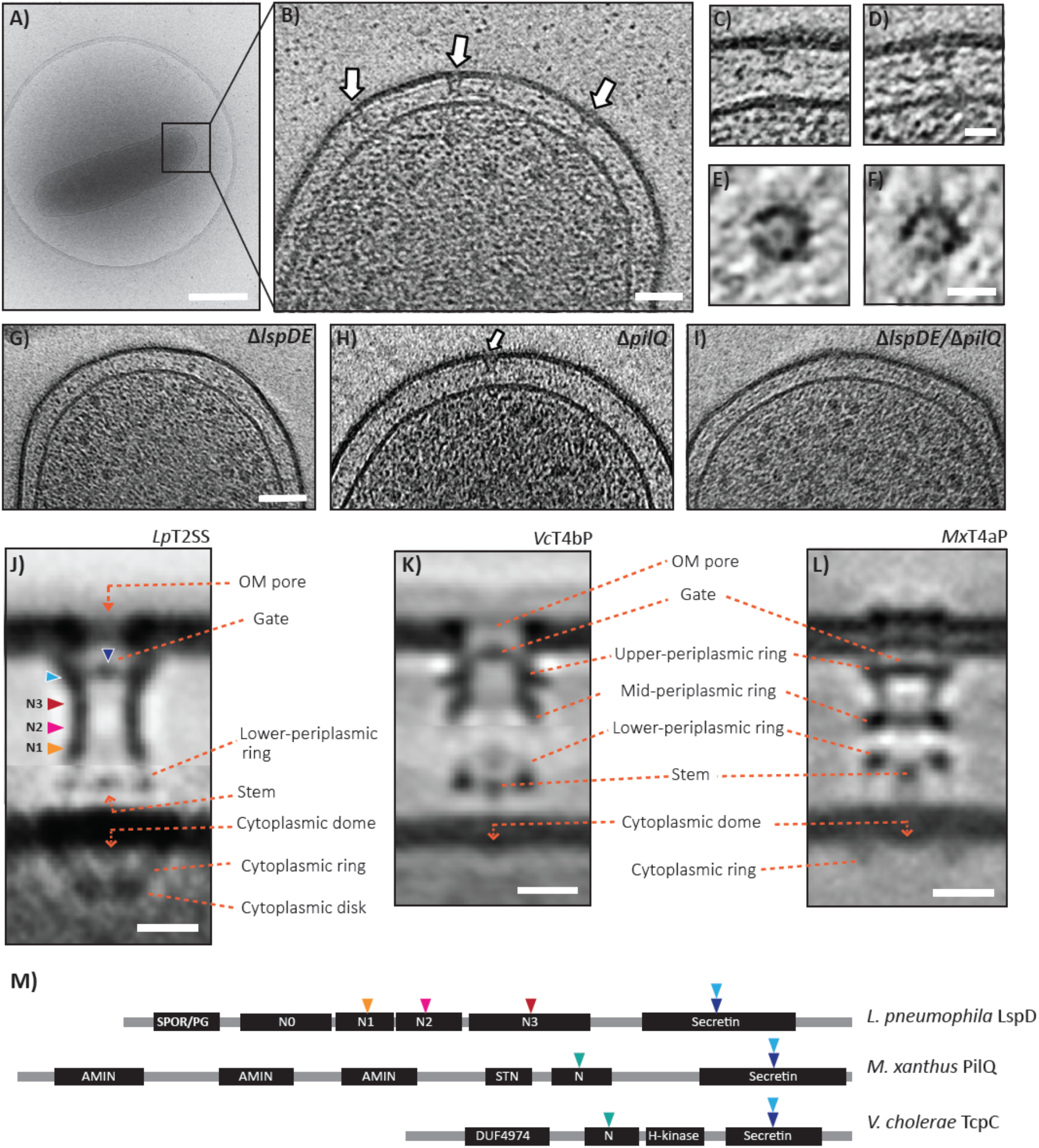
Visualization of the type II secretion system (T2SS) in frozen-hydrated *L. pneumophila* cells. (A) Cryo-EM image of a frozen-hydrated *L. pneumophila* cell on a Quantifoil grid. (B) Central slice through a tomographic reconstruction of a *L. pneumophila* cell pole. White arrows point to “hour-glass” shaped putative T2SS structures. (C and D) Tomographic slices showing individual T2SS particles. (E and F) Top views of individual T2SS particles. (G-I) Tomographic slices of *L. pneumophila* mutants lacking the T2SS secretin (G), the type IVa pili secretin (H), or both (I). (J) Composite subtomogram average of the wild-type *L. pneumophila* T2SS. (K-L) Subtomogram averages of the *Vibrio cholerae* T4bP and *Myxococcus xanthus* T4aP^39,40^ for comparison. (M) Domain architectures of the secretin protein in the *L. pneumophila* T2SS (T2S D or LspD), *M. xanthus* T4aP (PilQ) and *V. cholerae* T4bP (TcpC). Protein regions corresponding to different densities are indicated by arrowheads. PG: Peptidoglycan binding domain, SPOR: Sporulation related repeat, STN: Secretin and TonB N terminus short domain. Scale bars, 500 nm (A), 50 nm (B), 20 nm (C, D), 10 nm (E, F), 50 nm (G-I), 10 nm (J-L).

To reveal the *in situ* molecular architecture of the intact *L. pneumophila* T2SS, we generated a subtomogram average using 440 particles. In our initial average, the OM and secretin channel resolved well, but the IM and associated structures were missing (Supplementary Fig. 1A), suggesting substantial flexibility between the OM-associated secretin complex and the rest. To overcome this, we performed focused alignment on the OM- and IM-associated subcomplexes separately and then combined the two well-aligned averages to produce our final composite structure (Fig. 1J, Supplementary Fig. 1A-C) which had a variable local resolution between ~2.5-4.5 nm as estimated by ResMap (Supplementary Fig. 1D). The final average revealed that the T2SS is composed of a ~23 nm long vestibule that reaches down most of the way through the periplasm and opens up in the OM with an ~8 nm wide pore (Fig. 1J). There are two distinct densities in the lumen of the secretin channel near the two ends: 1) a “gate” just underneath the OM pore and 2) a previously unreported structure, “plug”, near the base (Supplementary Fig. 1E). Although the gate structure resolved well in our subtomogram average, the plug structure was poorly visible. Further analysis revealed that the plug is either not present or so dynamic in a subset of the particles that it is almost invisible in the final average (Supplementary Fig. 1E). Below the secretin channel, 7.5 away from the IM, there is a lower-periplasmic ring and a short stem at the same level (Fig. 1J). The lower-periplasmic ring is 17 nm wide in diameter (peak to peak) and the stem structure is in the middle of this ring (Fig. 1J). On the cytoplasmic side, there are three distinct densities: a dome-like density immediately below the IM, a ring-like density of diameter 20 nm (peak to peak) (cytoplasmic ring) surrounding the dome and finally, 13 nm away from the IM a 12-nm wide (peak to peak) cytoplasmic disk (Fig. 1J). A comparison between the subtomogram averages of the T2SS and the non-piliated *in situ* structures of the *Myxococcus xanthus* T4aP (*Mx*T4aP) and the *Vibrio cholerae* toxin-coregulated (TCP) type IVb pilus (VcT4bP)^39,40^ machines revealed that despite varied gene organization and limited component homology, the overall architectures of these three classes of molecular machines are strikingly similar: Each contains an OM-associated secretin channel, a gate just below the OM, a lower-periplasmic ring near the IM, a stem, a cytoplasmic dome, a cytoplasmic ring and a cytoplasmic disk (Fig. 1 J-L, note cytoplasmic disks for the *Mx*T4aP and *Vc*T4bP are seen in other states. See Supplementary Fig. 1F).

Intriguingly, the *L. pneumophila* secretin (T2S D, aa = 791) sequence is shorter than the *Mx*T4aP secretin (PilQ, aa = 901), but in the subtomogram averages, the *L. pneumophila* T2SS secretin channel is ~1.5 times longer than the *Mx*T4aP. To rationalize this, we predicted domain architectures of secretins of the *L. pneumophila* T2SS (T2 D or LspD), *Vc*T4bP (TcpC) and *Mx*T4aP (PilQ) systems using Motif-search, Phyre and the CD-vist programs^41,42^ (Fig. 1M). Our analysis revealed that all three secretins (*Lp*T2 D, *Mx*PilQ and *Vc*TcpC) have conserved C-terminal secretin domains. However, preceding the secretin domain, TcpC and PilQ have only one N-domain, but T2S D has four (N3, N2, N1 and N0) (Fig. 1M, see the match of the secretin atomic models to the subtomogram average density described below and shown in Supplementary Fig. 2). N-domains are known to fold into rigid structures and thus resolved well in the subtomogram average^39,43^. In contrast, the N-terminal AMIN domains of PilQ spread on the peptidoglycan (PG) and are not ordered and thus remained indiscernible in the subtomogram average.

In our subtomogram average, we saw densities for the cytoplasmic dome, ring and disk of the T2SS (Fig. 1J). The cytoplasmic dome is also visible in the piliated as well as in the non-piliated state of the *Mx*T4aP and *Vc*T4bP machines (Fig. 1K,L, Supplementary Fig. 1F), but the cytoplasmic ring and the disk are only visible in certain states of the *Mx*T4aP and *Vc*TcpC systems interpreted previously as assembly states (Fig. 1J-L, Supplementary Fig. 1F).

Our *in situ* structure provides the first glimpse of how the T2SS is positioned in the cell envelope with respect to the OM, PG and the IM. In our average, the secretin complex forms an OM-spanning pore. However, when we aligned available cryoEM structures of assembled secretin channels onto our density map based on the position of the gate (Supplementary Fig. 2), the tip of the secretin penetrated only the inner leaflet of the OM, suggesting secretin assemblies might have a different, more extended conformation *in vivo*, and collapse or change conformation during detergent extraction and purification.

To investigate if the T2SS is activated under the growth conditions used, we looked for T2SS-effector release in the *L. pneumophila* culture supernatant. Western blot analysis of the *L. pneumophila* culture supernatant showed consistent presence of the effectors CelA, ProA and LegP during mid-log phase and early stationary phase, suggesting that at least some of the T2SSs are active (Fig. 2P). Release of these effectors is T2SS-specific because a T2SS-nonfunctional mutant *(ΔlspDE)* did not release any of these effectors (Fig. 2P, lane 4). Interestingly, in a strain lacking the functional Dot/Icm type IV secretion system (T4SS), we found 25% more T2SS particles. However, this does not correlate with an increase in T2SS effector release, since both a mutant deleted for all *dot/icm* T4SS genes and a mutant lacking *dotH, dotG* and *dotF* did not exhibit consistent increases in T2SS effector release over many experiments (one experiment shown in Fig. 2P, lanes 2, 3). Since we see T2SS-effector release in the culture media, at least some of the T2SS particles are in actively-secreting states.

**Fig. 2.**
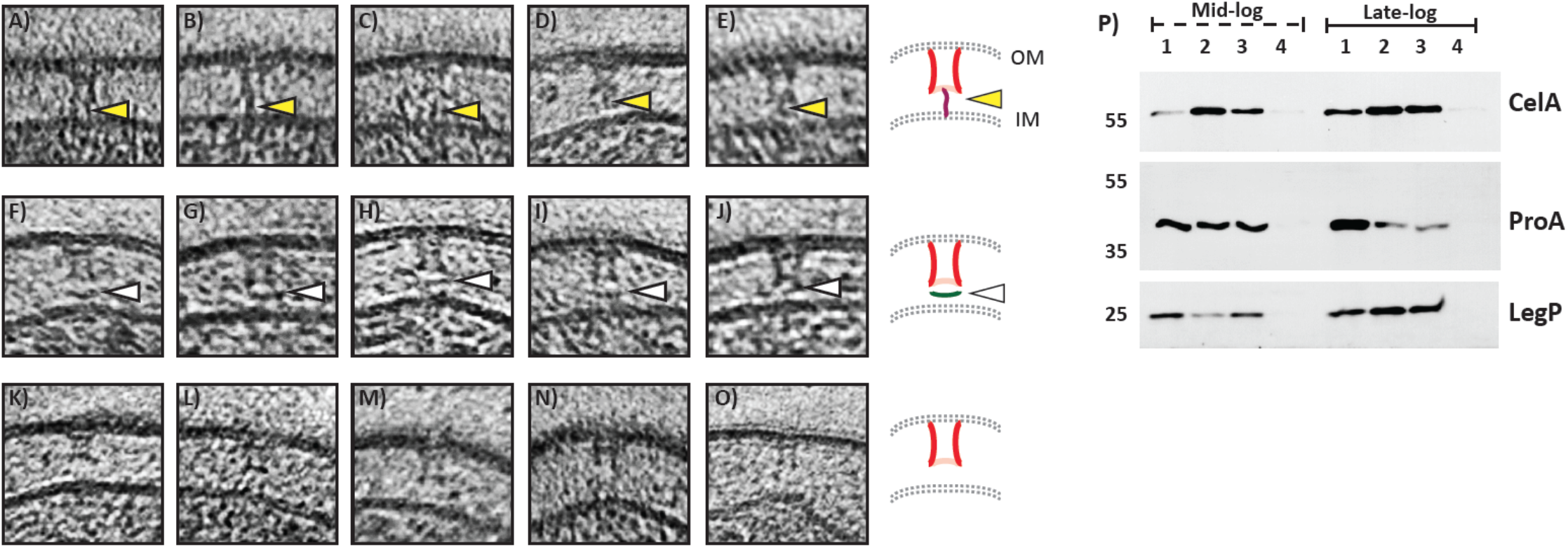
Individual T2SS subcomplexes. Tomographic slices showing individual T2SS particles with: (A-E) pseudopili (yellow arrowheads), (F-J) lower-periplasmic rings (white arrowheads) and (K-O) only secretin channels. Schematic representations (right). (P) Western blot analyses of T2SS-effectors (CelA, ProA and LegP) present in *L. pneumophila* culture supernatants at mid- and late-log phases. Different strains were analyzed: lane-1) WT *L. pneumophila* (Lp02), and control strains lacking 2) all the components of the *L. pneumophila* Dot/Icm T4BSS (JV4044), 3) a functional Dot/Icm T4BSS (ΔdotH, *ΔdotG*, and *ΔdotF:* JV7058) and 4) a functional T2SS *(ΔlspDE).* In several rounds of experiments, we did not notice any consistent increase in T2SS effector release in Dot/Icm T4SS nonfunctional mutants. Shown here are representative western blots.

In the T2SS, four minor pseudopilins (T2S I, J, K and H) and a major pseudopilin (T2S G) constitute a pseudopilus that is thought to act as a piston to extrude exo-proteins through the T2SS^6,16,17^. In mutant strains lacking the pseudopilins, the activity of the T2SS is severely impaired^44^. Since we did not see a pseudopilus in our subtomogram average (Fig. 1J), we carefully looked through all the individual T2SS particles in our tomograms and found that a fraction (~20%) of the particles had density likely from pseudopili (Fig. 2A-E). In these particles, the putative pseudopili extend (~ 14 nm) from the IM to the base of the T2S D channel. The positions of these densities are variable, however, so they remain invisible in the average. During this exercise, we also noticed that the lower-periplasmic ring densities in individual particles were just as visible as the secretin density, but their positions varied between particles, explaining why this density also resolved poorly in the subtomogram average (Fig. 2F-J). Further, there were also many particles with just the periplasmic vestibule and without a detectable pseudopilus or a lower-periplasmic ring (Fig. 2K-O). These particles are likely either assembly or disassembly intermediates.

### Architectural model of an intact T2SS

Recently, Chang *et al* used a combination of ECT, subtomogram averaging and genetic manipulations to determine the locations of all the major components in the *Mx*T4aP and VcT4bP systems and produced an architectural model of the *Mx*T4aP machinery^39,40^. In order to assign the locations of T2SS components in our density map and build an architectural model of the T2SS, we performed a thorough sequence analysis and confirmed that the *L. pneumophila* T2SS components T2S C, D, E, F, G, and M are homologues of the T4aP components PilP, PilQ, PilB, PilC, PilA and PilO respectively^45^. Interestingly, the cytoplasmic domain of T2S L is homologous to the T4aP component PilM (Phyre: 30% coverage, 97% confidence and 15% identity) and the periplasmic domain of T2S L is homologous to the T4aP component PilN (Phyre: 23% coverage, 78.6% confidence and 13% identity) suggesting T2S L is a fusion of the T4aP proteins PilM and PilN.

Since atomic models are available for all the soluble domains of the T2SS components^14,15^, we sought to build an architectural model of the intact T2SS guided by the T4aP model (Fig. 3). The *L. pneumophila* T2SS has a total of 12 components. We began by placing an atomic model of the OM protein T2S D/secretin (PDB ID: 5WQ8) in our density map. The position of T2S D along its axis (perpendicular to the cell envelope) was set by aligning the gate densities. The N1, N2 and N3 domains of 5WQ8 matched well with the subtomogram average density map (Supplementary Fig. 2). While the N0 domain of secretin was not resolved in most of the previously reported single particle reconstructions, presumably due to flexibility, in our subtomogram average we saw an additional density for the N0 domain just below N1. Our interpretation is that the N0 domain is stabilized by the presence of its connecting subcomplexes and the rest of the cellular envelope. A co-crystal structure of the T2S D N0 domain and the C-terminal homology region (HR) domain of the clamp protein T2S C (PDB ID: 3OSS) was therefore placed in this density below the N1 domain. Since no oligomeric structure is available for T2S C, we used known information about symmetry, connectivities and orientation (e.g. towards membrane etc.) to first generate numerous 15-mer (the known symmetry of T2S D) ring models for 3OSS using SymmDock^46^ and placed the ring model in the electron density map that fit the density best and satisfied all other criteria. Recently, Chernyatina *et al* showed that *Klebsiella pneumoniae* T2S DLME constitute a 15:6:6:6 complex and 6-12 copies of T2S C are bound to this assembly^21^. Therefore, we removed 9 copies of T2S C randomly around the ring in the model to reflect this lowest stoichiometry and emphasize that not all T2S D’s are bound by a T2S C^21^. Other than the C-terminal HR domain, *L. pneumophila* T2S C contains a periplasmic disordered region and an N-terminal transmembrane helix. We modeled the disordered region of T2S C and placed the N-terminal helix in the inner membrane.

**Fig. 3.**
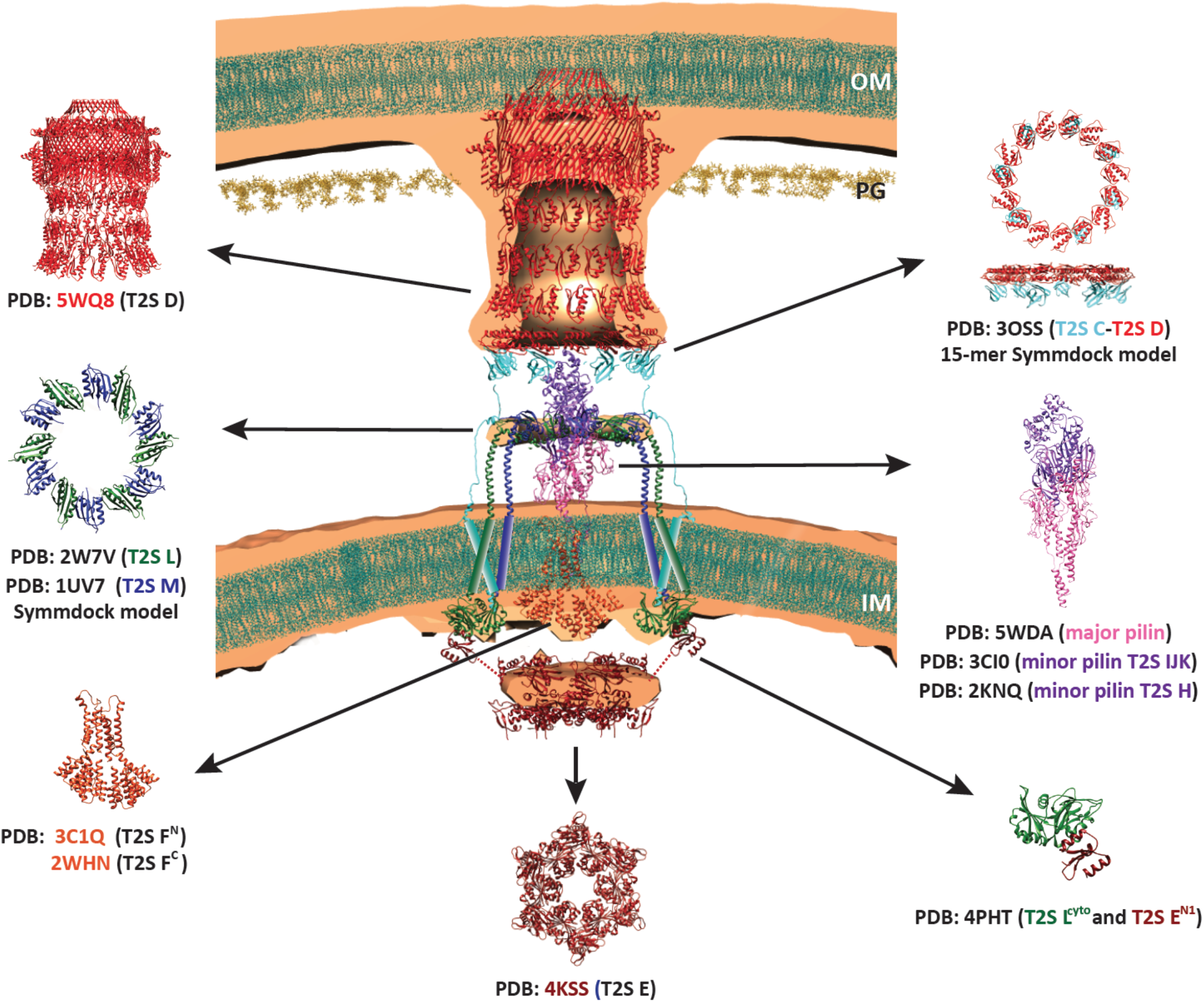
Architectural model of the T2SS. Atomic models of T2SS-components are superimposed on the central slice of the T2SS subtomogram average based on known connectivities and interfaces (see text). Transmembrane domains of the IM proteins are shown as cylinders. Linkers between N1E domain and N2E/CTE domains of T2S E are represented by dotted lines. OM = outer membrane, IM = inner membrane, PG = peptidoglycan. Lipids are shown in dark cyan and peptidoglycan dark yellow.

The periplasmic domains of T2S L (PDB ID: 2W7V) and M (PDB ID: 1UV7) both fold into ferredoxin-like domains and are known to bind each other^47^. We used a T2S M dimer structure (PDB ID: 1UV7) to first generate a T2S ML heterodimer and then used SymmDock to produce several hexameric ring models of the T2S ML heterodimer. We selected the model that best matched the lower-periplasmic ring-density. To build a working model for the pseudopilus, we used a cryoEM structure of the major pseudopilin filament (PDB ID: 5WDA), a co-crystal structure of the minor pseudopilin complex T2S IJK (PDB ID: 3CI0) and a crystal structure of the minor pseudopilin T2S H (PDB ID: 2KNQ) and their known connectivities and interfaces^26,27^. Since in our tomograms we only found pseudopili extending from the IM to the base of the secretin channel, we found that ~5 copies of major pseudopilin (T2S G) and one copy of each of the minor pseudopilins (T2S I, J, K, H) were sufficient to extend this distance.

To begin to generate a working model for the cytoplasmic complex, we modelled a T2S F dimer structure based on the proposed structure of the T4aP homologue PilC^40^ and placed this model in the cytoplasmic dome density. The T2SS ATPase T2S E has three distinct domains (N-terminal domains N1E, N2E and a C-terminal ATPase domain CTE) connected by two extended flexible linkers. We used a hexameric structure of the N2E+CTE domains of T2S E (PDB ID: 4KSS) and a co-crystal structure of the cytoplasmic domain of T2S L and the N1E domain of T2S E (PDB ID: 4PHT) to build working models for the cytoplasmic disk and ring densities, respectively. The cytoplasmic ring is 20 nm wide in diameter in our average. Given the experimentally determined stoichiometry of six T2S L molecules per secretin channel, we first used SymmDock to generate candidate ring assemblies of six 4PHT complexes. Interestingly, out of 5000 candidate ring models, none had a diameter more than 17 nm. Our interpretation of this result is that the T2S L:N1E complex does not form a continuous ring *in situ.* Rather we simply placed six separate copies of 4PHT around the cytoplasmic ring with gaps in between. The 4KSS structure fits nicely into the cytoplasmic disk as expected. Finally, T2S O is an IM peptidase that processes the pseudopilins prior to their assembly. This protein has no significant domain outside the membrane and is not visible in our subtomogram average. Therefore, we did not place this protein in our architectural model.

## Discussion

Here, we present the first structure of an intact T2SS *in situ.* Compared to the *Mx*T4aP and the *Vc*T4bP systems, the basic architecture of these molecular machines is clearly very similar (Fig. 1J-L, Table 1). Since form follows function in biology, this points to overall conservation of basic function (assembling and disassembling a pilus that grows from the IM towards or through the OM) despite the presence of components without recognizable sequence homology. There are also many interesting differences, however, which lead to specific hypotheses about the roles and adaptations of many of the components (Table 1).

**Table 1:**
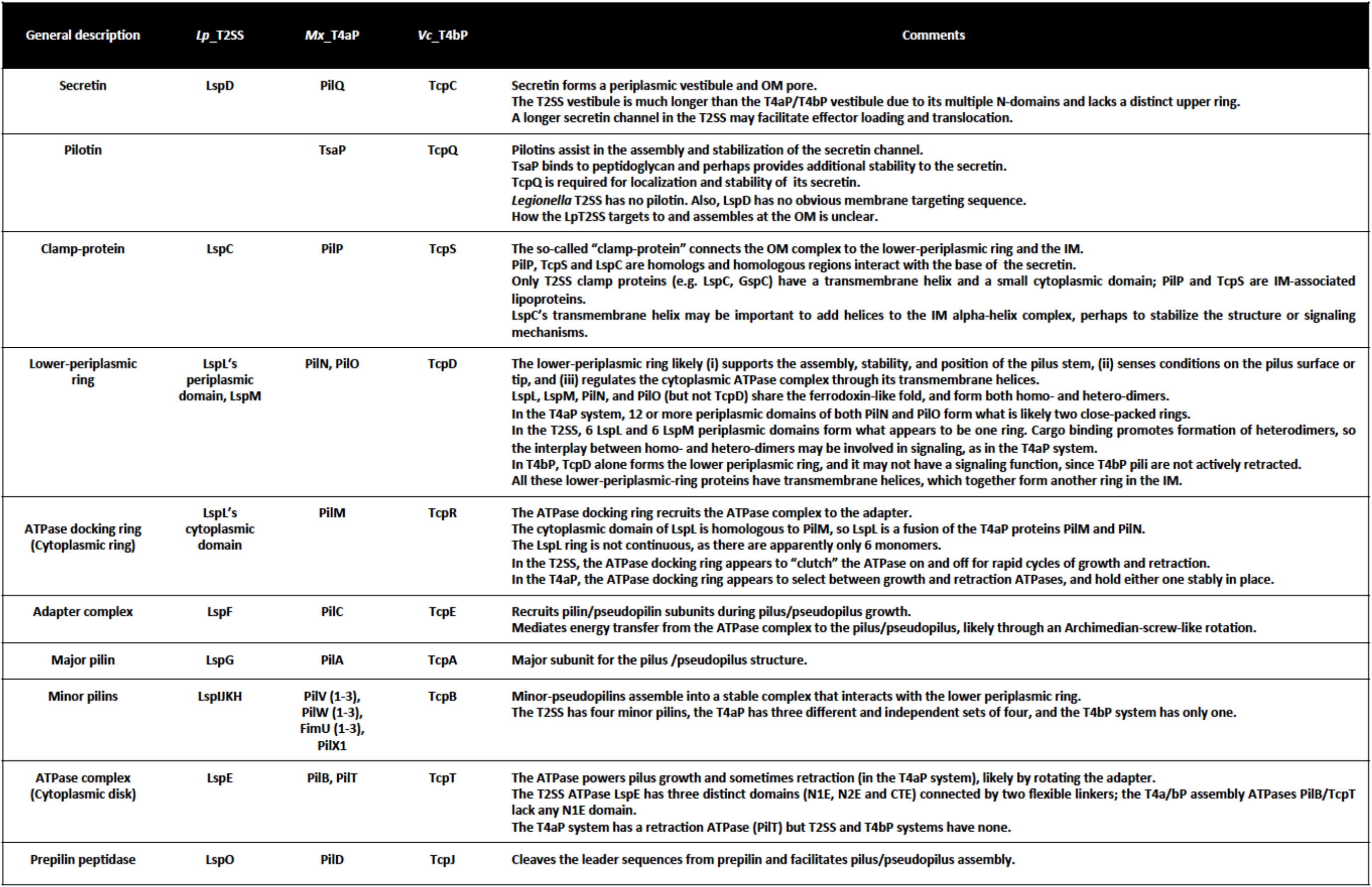
Comparison between the T2SS and related molecular machines T4aP and T4bP systems.

First, the T2SS has a much longer periplasmic vestibule (23 nm) than the T4aP and T4bP systems (15 nm). The T2SS translocates several exoproteins through the periplasmic secretin channel, whereas the T4aP and T4bP systems secrete only their pili. We speculate that the T2SS secretin evolved a large vestibule to load cargo. Consistent with this notion, in our *in situ* structure, we observed a poorly-resolved plug density near the base of the secretin vestibule, close to the N1 domain of T2S D (Supplementary Fig. 1C). T2S C is known to interact with the N0 domain of T2S D through its HR domain and recruit T2SS effectors into the complex^25^. The plug density could be the HR domain of T2S C^25^ or cargo that is waiting to be extruded by the pseudopilus.

Unlike the *Mx*T4aP and the VcT4bP systems, the *L. pneumophila* T2SS does not encode a pilotin^48^, consistent with the idea that pilotins are not essential for all secretin pores to assemble. It remains unclear how the *L. pneumophila* T2 D is targeted to the cell pole and how it associates with the PG. In the *Mx*T4aP system, the N-terminal AMIN domains of PilQ tether the T4aP machine to the PG layer^40^. In our *L. pneumophila* tomograms, the PG layer is located immediately beneath the OM (Supplementary Fig. 3) but the putative PG binding domain of T2S D is N-terminal to the N0 domain, which is ~20 nm below the PG (Fig. 1M). One possibility is that the linker between the N0 domain and the putative PG binding domain bridges this gap. Curiously, the *L. pneumophila* T2SS also lacks other PG binding complexes like those present in *Vibrio spp, Aeromonas spp* and in *E. coli* T2SS (e.g. *gspAB* homologs)^49^.

Here, we found that the periplasmic domains of T2S L and M form a lower-periplasmic ring similar to those present in the *Mx*T4aP and the VcT4bP systems. PilO and PilN (the two proteins that form the lower-periplasmic ring in the *Mx*T4aP), and T2S L and T2S M all begin with an N-terminal transmembrane helix, followed by a ferrodoxin-like fold in the periplasm. Chang *et al*. previously hypothesized that in the *Mx*T4aP, the lower-periplasmic ring likely transmits information about the status of the pilus to the cytoplasmic subcomplex^40^. Given that T2S L and T2S M have recently been shown to interact with secreted effectors^47^, T2S L and M may likewise sense cargo in the periplasm and signal the cytoplasmic complex to extend the pseudopilus. Signal sensing and transmission are likely simpler in the T2SS, however, since while the lower-periplasmic ring in the *Mx*T4aP is composed of 12 or more copies of both PilO and PilN, in *L. pneumophila*, the lower-periplasmic ring is apparently comprised of only six copies each of T2S L and M. Based on these stoichiometries, the conservation of fold, and the appearances of the rings in the sub-tomogram averages, we predict that the basic structures of these lower-periplasmic rings are the same, but there are two closely-packed rings in the *Mx*T4aP, as opposed to only one in the T2SS. Signaling likely involve transitions between homodimer and heterodimer associations, as the presence of cargo biases the interactions between T2S L and M towards heterodimers^47^, and the *Vc*T4bP system, which may not signal (the pilus is never actively retracted by an ATPase), has only a single lower-periplasmic-ring protein (TcpD)^39^. It remains elusive how the symmetry mismatch between the secretin channel, T2S C and the lower-periplasmic ring is coordinated and what the role of symmetry mismatch is in effector loading and export.

The conservation of transmembrane helices in all the lower-periplasmic-ring proteins suggests an important role in signal transmission or structure, but the number of helices present and their connections to the cytoplasmic ring vary. Assuming there are six copies of both T2S L and M, a ring of 12 transmembrane helices will be present in the IM. This contrasts with the 24 or more helices from PilN and PilO in the *Mx*T4aP. Perhaps in partial compensation, the T2SS clamp protein T2S C begins with a transmembrane helix, adding 6-10 more helices to the ring (according to current estimates of stoichiometry), while its counterparts in the *Mx*T4aP and *Vc*T4bP systems (PilP and TcpS, respectively) are simply lipoproteins, and so add nothing. The consequence of this is not immediately clear, but it is known that T2S C’s oligomerization and interaction with T2S L/M are regulated through the T2S C transmembrane helix^50^, and that the transmembrane regions of T2S L, M and C form a dynamic network^15,50^. Curiously, in the *Vc*T4bP system, there are again two proteins contributing alpha helices in the IM, but in this case one is from the periplasmic ring (TcpD) and the other is from the cytoplasmic ring (TcpR). The *L. pneumophila* T2S exhibits yet another variation, in that one lower-periplasmic-ring protein (T2S L) also forms part of the cytoplasmic ring, with the transmembrane helix in between. We speculate that the fusion allows another adaptation, which is that in *L. pneumophila* the cytoplasmic domain of T2S L appears to not form a continuous ring. Being fused to the periplasmic ring may be important to hold it in place. From all this, it appears that there is an important network of 18 or more interacting transmembrane helices in the IM that mediate signal transmission between the lower-periplasmic ring and the cytoplasmic ring.

Here, we found that just as in the *Mx*T4aP and the *Vc*T4bP systems, the T2SS ATPase sits directly below the IM on the machine’s axis. The main difference is, however, that it is positioned further away from the IM and is less well resolved. We speculate that as the T2SS rapidly switches from extending to retracting the pseudopilus (the piston model, see below), it does so by moving the ATPase up against the adaptor T2S F to extend the pseudopilus, and then dropping it away from T2S F to let the pseudopilus retract. Our interpretation of the data is, then that most of our particles were in the retraction state, with the ATPase disengaged from T2S F. This mechanism is of course different than the *Mx*T4aP system, which has two ATPases, one responsible for extension and the other retraction^40^. In the *Mx*T4aP case, the two ATPases occupy the same position (they are exchanged) and were seen in contact with the adaptor in the cryotomograms, suggesting they are held there continuously for long periods of time. In contrast, the T2SS likely evolved to rapidly switch between short bursts of extension and retraction, and so it does not release its ATPase completely – it appears to merely “clutch” it on and off. In the retraction state, we propose that the N1E domain is close to the IM and intesracts with the cytoplasmic domains of T2S L and T2S F^24,51–53^, while the N2E and CTE domains dangle on flexible linkers, causing them to resolve poorly in our average.

Several individual particles exhibit putative pseudopili that extend from the IM to the base of the T2S D channel. No pseudopilus was seen to extend up to the gate, suggesting either the pseudopilus only pushes exo-proteins into the periplasmic vestibule or an extended form of a pseudopilus (reaching up to the gate or beyond) is short-lived. Our model showed that only ~5 copies of the major pseudopilin T2S G and one copy of each of the minor pseudopilins (T2S I, J, K, H) is sufficient to span the distance between the IM and the base of the secretin. We also found that ~12-15 copies of the major pseudopilin and one copy of each of the minor pseudopilins T2S I, J, K, H would be required for the pseudopilus to extend to the secretin gate. It is not clear how the length of the pseudopilus is controlled. Intriguingly, the C-terminal domain of T2S L has sequence similarity (Phyre: 52 amino acids, 68% confidence and 15% identity) to the C-terminal domain of FliK that controls flagellar hook length^54^. Thus, T2S L may have multiple functions.

The piston model for substrate extrusion posits that the T2SS iteratively extends and retracts a short but permanent pseudopilus, pushing cargo out of the cell^55^. An alternative model is that the pseudopilus completely disassembles after cargo release, and a completely new pseudopilus assembles in the next round^17^. Concerning these possibilities, we note that the stem density positioned right in the middle of the lower-periplasmic ring likely represents the minor pseudopilin complex. Because this stem density is clear in the sub-tomogram average, it means it is usually present. Second, the cytoplasmic disk is the ATPase, revealing that it too is usually present. Both points support the piston model.

Overall, our *in situ* structure of the *L. pneumophila* T2SS highlights commonalities and key differences between the evolutionarily-related T2SS, T4aP and T4bP machines and provides new insights into its structure and function.

## Materials and Methods

### Strains, growth conditions, and mutant generation

All experiments were performed using the *L. pneumophila* Lp02 strain (thyA hsdR rpsL), a derivative of the clinical isolate *L. pneumophila* Philadelphia-1. Cells were grown as described previously^57,58^. Briefly, cells were grown in ACES buffered yeast extract (AYE) broth or on buffered charcoal yeast extract (CYE) plates. The culture media were always supplemented with thymidine (100 μg/ml), ferric nitrate and cysteine hydrochloride.

A T4SS mutant of Lp02 that lacks all 26 of the *dot/icm* genes (i.e., JV4044) was previously described^59,60^, as was a mutant of Lp02 that lacks *dotH, dotG*, and *dotF* (i.e., JV7058)^60^. To generate a *lspDE* mutant lacking the T2SS (i.e., strain NU438) and a *pilQ* mutant lacking T4P (i.e., strain NU439) of *L. pneumophila*, previously reported plasmids were introduced into Lp02 by natural transformation and then the desired mutations were introduced into the bacterial chromosome via allelic exchange^32^. Plasmid pOE4Kan was used to make the *lspDE* mutant, and pGQ::Gm was employed for making the *pilQ* mutant^61,62^. In a similar way, a *lspDE pilQ* double mutant lacking both T4SS and T4P (i.e., strain NU440) was constructed by introducing pGQ::Gm into strain NU438.

### Assay for secreted proteins

*L. pneumophila* strains were grown in AYE broth at 37°C to either the mid-log or late-log phase of growth, at which times the cultures were centrifuged and the resultant supernatants obtained and filtered through 0.2 μm syringe filters. Five μl of the supernatant samples were subjected to SDS-PAGE and then analyzed by immunoblot as previously described^37^. Briefly, after blocking in 5% milk for 1 h, the blots were incubated overnight with 1:1,000 dilutions of rabbit anti-CelA, anti-ProA, or anti-LegP antibodies, washed, and then incubated for 1 h with 1:1,000 dilution of secondary HRP-conjugated goat anti-rabbit IgG antibody (Cell Signaling Technology). Images of the immunoblots were developed with ECL^TM^ Western Blotting Detection Reagent (GE Healthcare).

### Sample preparation for electron cryotomography

*L. pneumophila* Lp02 cells were grown till early stationary stage (OD_600_ ~2.8) and harvested. Cells were mixed with 10-nm colloidal gold beads (Sigma-Aldrich, St. Louis, MO) precoated with bovine serum albumin. Four μl of this mixture was applied onto freshly glow-discharged copper R2/2 of th0 Quantifoil holey carbon grids (Quantifoil Micro Tools GmbH, Jena, Germany). Using an FEI Vitrobot Mark IV, grids were then blotted (under 100% humidity conditions) and plunge-frozen in a liquid ethane/propane mixture.

### Electron tomography and subtomogram averaging

The frozen grids were subsequently imaged in an FEI Polara 300 keV FEG transmission electron microscope (Thermo Fisher Scientific) coupled with a Gatan energy filter and a Gatan K2 Summit direct electron detector. Energy-filtered tilt series of cells were collected automatically from −60° to +60° at 1.5° intervals using the UCSF Tomography data collection software^63^ with a cumulative total dosage of 100 e^−^ Å^−2^, a defocus of −6 μm and a pixel size of 3.9 Å. Using the IMOD software package^64^, the images were then binned by 2, aligned and contrast transfer function corrected. Subsequently, SIRT reconstructions were produced using the TOMO3D program^65^. T2SS structures on cell envelopes were visually identified by their characteristic “hour-glass” like shape. Subtomogram averages of the *L. pneumophila* T2SS were generated by the PEET program^66^. The T2SS subtomogram averages exhibited a gross twofold symmetry around the central midline in the periplasm. Based on this observation, we applied a two-fold symmetry on the periplasmic complex. No symmetry was applied on the cytoplasmic complex. The T2SS exhibited a substantial flexibility between the OM- and IM-associated parts; binary masks were applied to perform focused alignments on the OM and on the IM complexes separately and finally combined to generate a composite structure. The numbers of tomograms collected and number of particles used are summarized in Supplementary Table 1.

### Identifying homologues between the *L. pneumophila* T2SS and the T4P system

*L. pneumophila* Lp02 strain T2SS protein sequences were selected from the NCBI and UniProt databases. To identify corresponding components between the *L. pneumophila* T2SS and the *Mx*T4aP and *Vc*T4bP systems, we utilized two different programs: Phyre and Blast search^41,67^. This confirmed *L. pneumophila* T2SS components T2S C, D, E, F, G and M are homologues of the T4aP components PilP, PilQ, PilB, PilC, PilA and PilO respectively. Using a combination of Phyre, MOTIF-search, and Blast programs, we also predicted distinct domains and motifs within different T2SS components.

## Data availability

The subtomogram average of the *L. pneumophila* T2SS has been deposited in the Electron Microscopy Data Bank under the following accession codes: XXXX (wild-type aligned on the OM part); EMD-XXX (wild-type, aligned on the IM part)

## Acknowledgements

This work was supported by NIH grants AI127401 to G.J.J. and AI043987 to N.P.C. ECT data were recorded at the Beckman Institute Resource Center for Transmission Electron Microcopy at Caltech and the cryo-EM facility at Janelia Research Campus. We thank Dr. Davi Ortega for helpful discussions. M. K. is supported by a postdoctoral Rubicon fellowship from De Nederlandse Organisatie voor Wetenschappelijk Onderzoek (NWO).

