## Supplementary material for "*In vivo* structure of the *Legionella* type II secretion system by electron cryotomography"

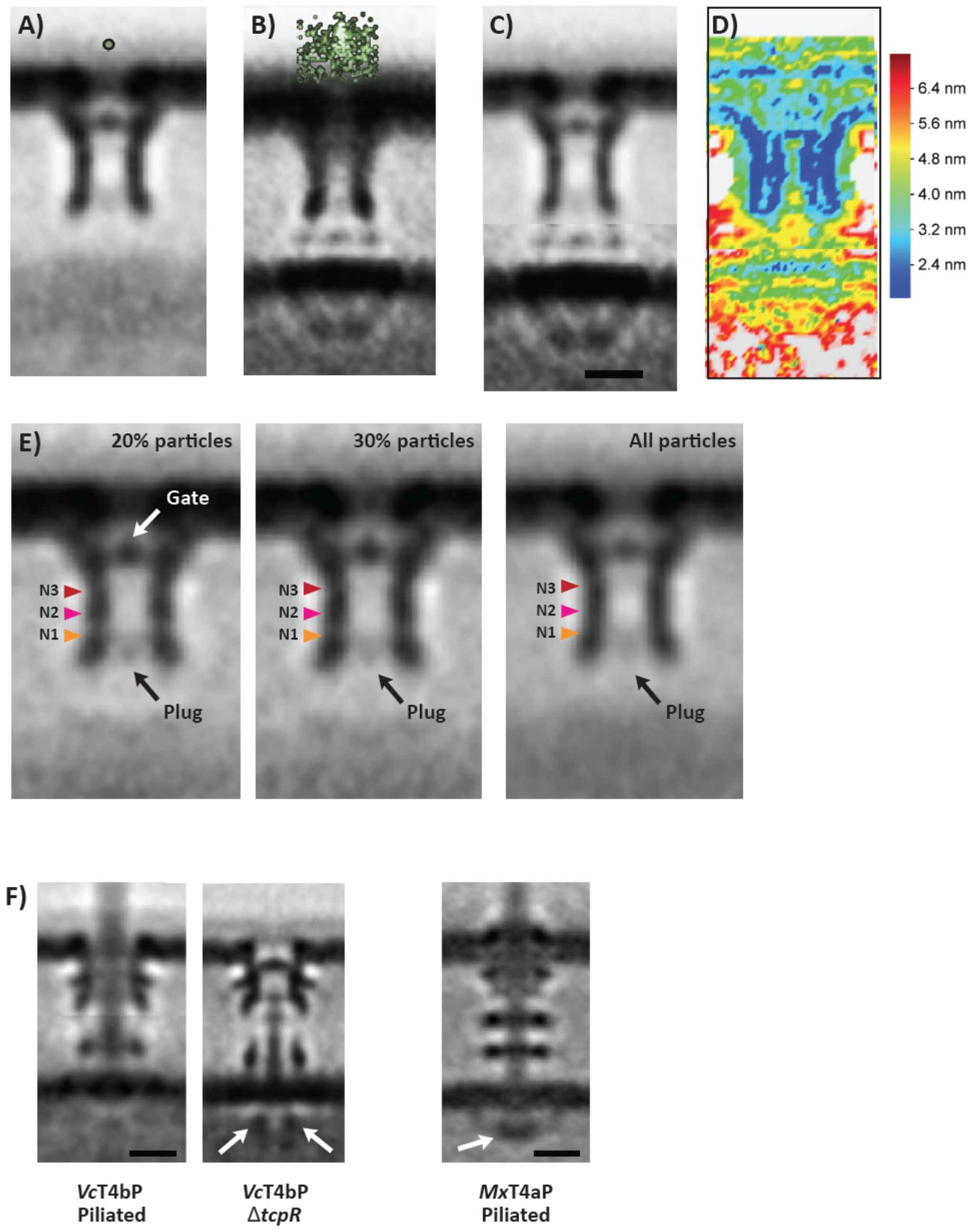


**Supplementary Figure 1. T2SS Flexibility.** Subtomogram average of all particles aligned on (A) the OM-associated complex and (B) the IM-associated complex. The distribution of the green dots in (B) indicates the translations imposed on the OM complexes to align the IM complexes. (C) A composite average using the upper and lower halves of (A) and (B), respectively. D) Local resolution of (C) calculated by Resmap[56]. (E) Focused alignment near the base of the secretin channel revealed the presence of a plug-like structure.

20% of the particles with highest cross-correlation showed this distinct density. In the rest of the particles, the plug density is either not present or so dynamic that including them makes the plug almost invisible. (F) Previously reported *in situ* averages of the T4aP (WT, piliated) and T4bP (WT, piliated and Δ*tcpR* mutant) machines in states with cytoplasmic dome, ring and disks for comparison[39,40]. White arrows indicate cytoplasmic disks.

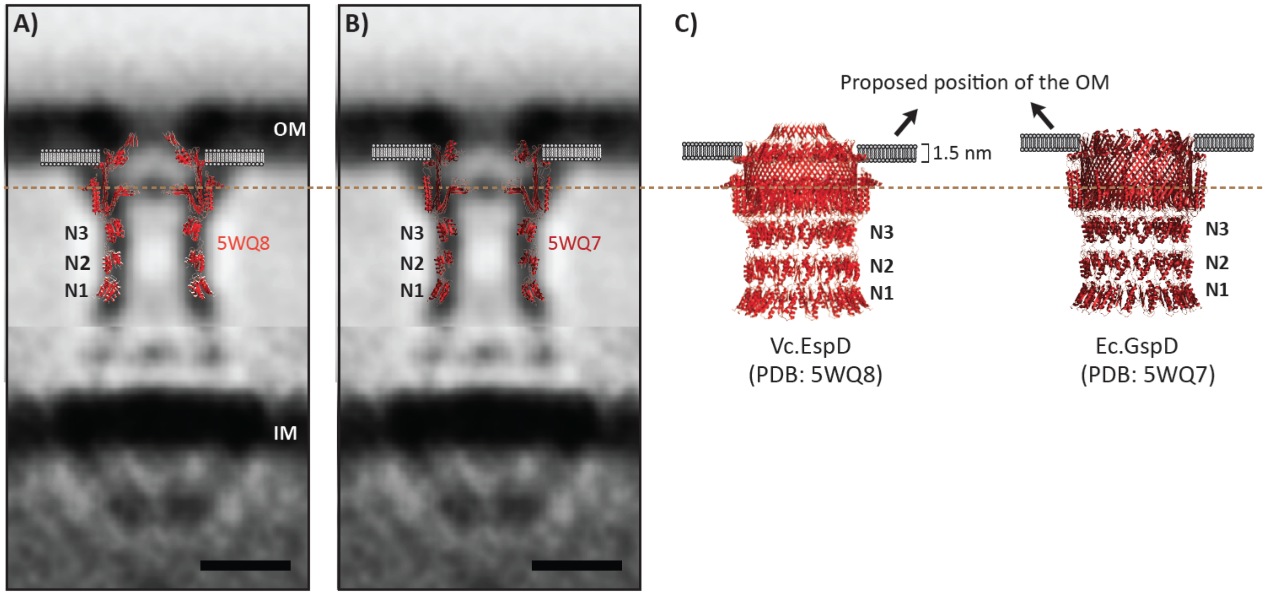


**Supplementary Figure 2. Position of the T2SS secretin with respect to the OM.**

(A and B) Atomic models of the *V. cholerae* (PDB ID: 5WQ8) and *E. coli* (PDB ID: 5WQ7) T2SS secretins superimposed on our subtomogram average based on the position of the gate. (C) Positions of the OM on these structures as suggested in earlier publications[18,20]. The widths of the suggested OM spanning regions were only ~1.8 nm, but real membranes are known to be 5-7 nm wide. In all reported atomic models, the secretin channel is suggested to extend beyond the OM[14,18]. However, when we overlaid the secretin atomic models on our subtomogram average, it only reached through the inner leaflet of the OM. Scale bars, 10 nm.

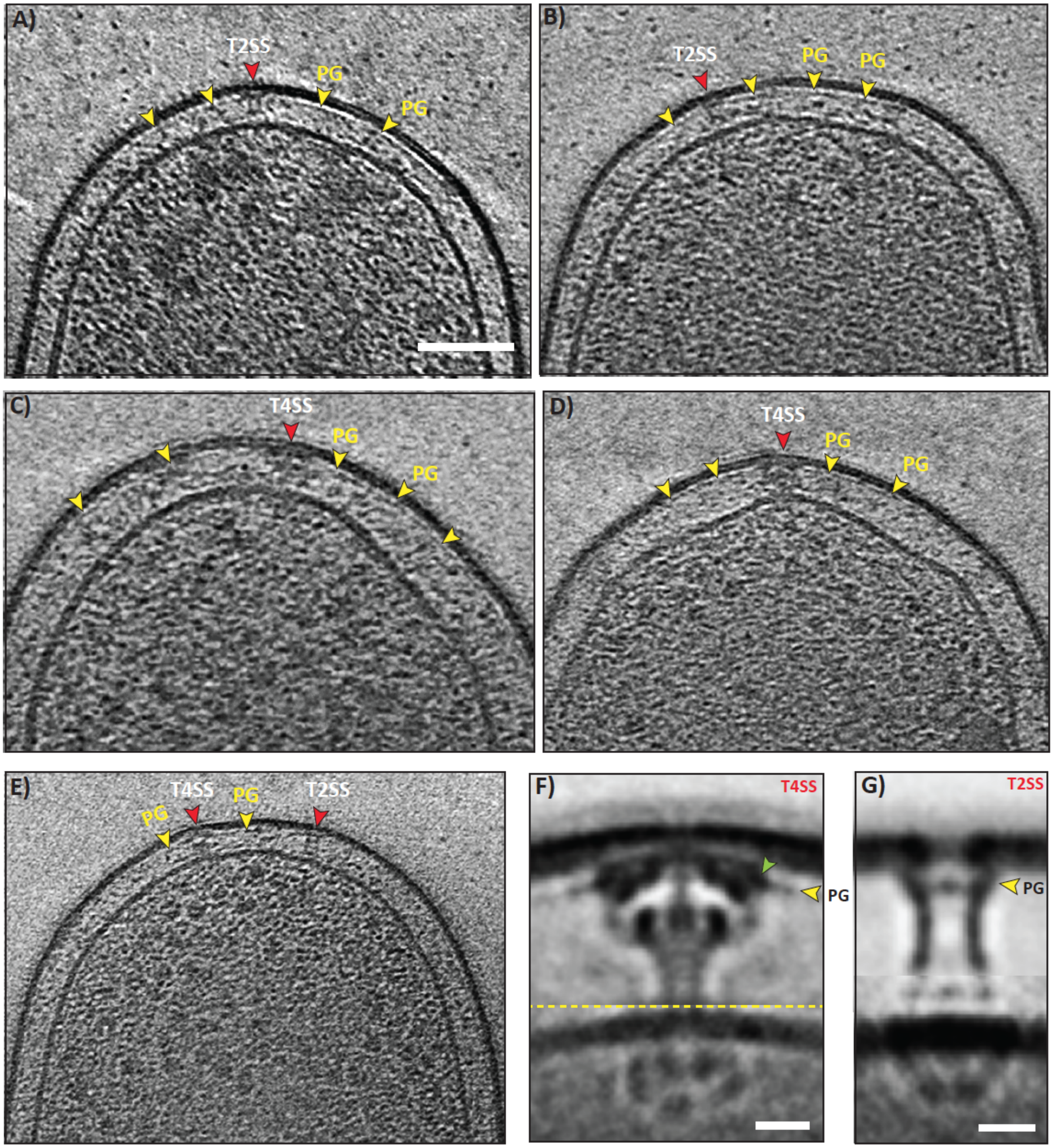


**Supplementary Figure 3. Position of the T2SS with respect to the PG layer in *L. pneumophila*.** (A and B) Tomographic slices through *L. pneumophila* cells showing T2SS particles (red arrowheads) and the peptidoglycan layer (PG, yellow arrowheads). (C and D) Tomographic slices through *L. pneumophila* cells showing T4BSS particles (red arrowheads) and the peptidoglycan layer (PG, yellow arrowheads). (E) Tomographic slice through a *L. pneumophila* cell showing both T4SS and T2SS particles (red arrowheads) and peptidoglycan (PG, yellow arrowheads) in the same cell. (F and G) Subtomogram averages of the T4SS and T2SS, respectively. DotK (shown as green arrow) in the T4BSS is known to interact with the PG layer confirming its location just a few nm below the OM (F). We therefore conclude that the PG layer surrounds the T2SS at approximately the level of the gate (G).

**Supplementary Table 1.**

| No | Strain name | Description | Tomograms collected | Particles found |
|---|---|---|---|---|
| 1 | Lp02 | *Legionella pneumophila thyA* Lp02 | 1993 | 440 |
| 2 | Lp02 (Δ*lspDE*) | *Legionella pneumophila thyA* Δ*lspDE* | 38 | None seen |
| 3 | Lp02 (Δ*pilQ*) | *Legionella pneumophila thyA* Δ*pilQ* | 41 | 14 |
| 4 | Lp02 (Δ*lspDE* /Δ*pilQ*) | *Legionella pneumophila thyA* Δ*lspDE* /Δ*pilQ* | 27 | None seen |
